# Traumatic Brain Injury Impairs Systemic Vascular Function Through Altered Lipid Metabolism and Disruption of Inward-Rectifier Potassium (Kir2.1) Channels

**DOI:** 10.1101/2021.01.15.426853

**Authors:** Adrian M. Sackheim, Nuria Villalba, Maria Sancho, Osama F. Harraz, Adrian D. Bonev, Angelo D’Alessandro, Travis Nemkov, Mark T. Nelson, Kalev Freeman

## Abstract

**BACKGROUND AND PURPOSE:** Trauma can lead to widespread vascular endothelial dysfunction, but the underlying mechanisms remain largely unknown. Strong inward-rectifier potassium channels (Kir2.1) play a critical role in the dynamic regulation of regional perfusion and blood flow. Kir2.1 channel activity is modulated by phosphatidylinositol 4,5-bisphosphate (PIP_2_), a minor membrane phospholipid that is degraded by phospholipase A_2_ (PLA_2_) in conditions of oxidative stress or severe inflammation. We hypothesized that PLA_2_–induced depletion of PIP_2_ impairs Kir2.1 channel function.

**METHODS:** A fluid percussion injury model of traumatic brain injury (TBI) in rats was used to study mesenteric resistance arteries 24 hours after injury. Patch-clamp electrophysiology in freshly isolated endothelial and smooth muscle cells was performed to monitor Kir2.1 conductance, and the functional responses of intact arteries were assessed using pressure myography. We analyzed circulating PLA_2_, hydrogen peroxide (H_2_O_2_), and metabolites to identify alterations in signaling pathways associated with PIP_2_ in TBI.

**RESULTS:** Electrophysiology analysis of endothelial and smooth muscle cells revealed a significant reduction of Ba^2+^-sensitive Kir2.1 currents after TBI. Additionally, dilations to elevated extracellular potassium and BaCl_2_- or ML 133-induced constrictions in pressurized arteries were significantly decreased following TBI, consistent with an impairment of Kir2.1 channel function. The addition of a PIP_2_ analog to the patch pipette successfully rescued endothelial Kir2.1 currents after TBI. Both H_2_O_2_ and PLA_2_ activity were increased after injury. Metabolomics analysis demonstrated altered lipid metabolism signaling pathways, including increased arachidonic acid, and fatty acid mobilization after TBI.

**CONCLUSIONS:** Our findings support a model in which increased H_2_O_2_-induced PLA_2_ activity after trauma hydrolyzes endothelial PIP_2_, resulting in impaired Kir2.1 channel function.

## Introduction

Traumatic injury represents the most common cause of death in individuals aged 1-44, with the majority of these fatalities due to traumatic brain injury (TBI).^1^ In addition to the primary mechanical injury, systemic inflammation subsequent to trauma contributes to coagulopathy, vascular leak and multi-organ dysfunction. Vascular endothelial cells (ECs) are especially vulnerable to damage from cellular debris and circulating factors such as damage-associated molecular patterns released into the bloodstream after an injury.^2 3 4 5^ Endothelial injury after severe trauma is an important independent predictor of coagulopathy, multi-organ dysfunction and death. Clinical studies have consistently demonstrated the existence of elevated biomarkers unique to the endotheliopathy of trauma (EoT) in distinct cohorts of trauma subjects, including pediatric trauma and traumatic brain injury, correlating with adverse outcomes.^6 7 8 9 10 11 12^ We previously studied mesenteric arteries in animals after a traumatic brain injury (TBI), and demonstrated impaired endothelial-dependent vasodilation occurs through a mechanism which involves uncoupling of endothelial nitric oxide synthase (eNOS).^13^ Recent advances in systems-level screening and analysis have demonstrated profound metabolopathies occur following hemorrhagic shock, traumatic brain injury, and burn injury, including pronounced alterations in lipid metabolism which would be expected to impact blood vessel and endothelial functions.^14 15 16 17 18 19^ Despite these recent advances, the fundamental, cellular mechanisms leading to alterations in systemic vascular function after trauma remain elusive. Here, we analyzed downstream consequences of TBI on a critical modulator of vascular function: the inward rectifier potassium channel Kir2.1.

There are 15 members of the inward rectifier potassium (Kir) channel family. The Kir2 family are strong inward rectifier K^+^ channels, which are activated by external K^+^ and require phosphatidylinositol 4,5-bisphosphate (PIP_2_) for activity. We and others have provided strong evidence that smooth muscle cells (SMCs) and ECs have Kir2.1 channels.^21 22 23 24 25 26^ Kir2.1 channel function plays a key role in setting arterial resting membrane potential, myogenic tone development, and cerebral blood flow regulation.^27 28^ We recently demonstrated impaired capillary EC Kir2.1 function in the cortical hemisphere contralateral to a brain injury; but prior studies have not addressed whether this dysfunction extends to blood vessels in the systemic circulation^29^. PIP_2_ is a minor membrane phospholipid degraded under conditions of oxidative stress or severe inflammation that affect lipid metabolism. Under these conditions, phospholipases (mainly from the A and C family) catalyze membrane phospholipids to form signaling molecules including inositol trisphosphate, diacylglycerol, and arachidonic acid (AA). Specifically, PIP_2_ is cleaved by PLA_2_ at the sn-2 acyl bond, freeing AA, which is subsequently modified by downstream cyclooxygenases and lipoxygenases into prostanoids and eicosanoids including prostaglandins and leukotrienes.^30^ Disturbed lipid metabolism in severely injured trauma patients is a strong predictor of clinical outcome.^31^ Altered fatty acid metabolism has been demonstrated in controlled animal models of stroke^32^ and tissue injury with hemorrhagic shock, including derangements in mono- and poly-unsaturated fatty acid mobilization and oxidation products.^17^ We hypothesized that severe TBI, even without concomitant shock or hypoxia, would cause significant metabolic responses. In this context, depletion of endothelial PIP_2_ in the plasma membrane may impair Kir2.1 channel function and therefore, vascular function. Here, we demonstrate a novel mechanism of altered systemic vascular function after TBI, through PIP_2_-dependent impairment of Kir2.1 channel function. Control of PIP_2_ levels may provide a therapeutic target to improve vascular function in conditions characterized by altered lipid metabolism such as TBI and stroke.

## Materials and Methods

The authors declare that all supporting data are available within the article (and its online supplementary files).

### Animal and injury model

Adult male Sprague-Dawley rats (aged 3 to 4 months; 300–325 □g; Charles River, Saint Constant, Quebec, Canada) were randomly assigned to either a fluid percussion TBI surgery or control treatment, as previously described.^33^ All procedures were approved by the Institutional Animal Care and Use Committee and were performed in accord with the National Research Council’s *Guide for the Care and Use of Laboratory Animals*. Expanded details are described in the Data Supplement.

### Sample Preparation and UHPLC-MS analysis

Plasma was collected and stored at −80°C until analysis. Analyses were performed as previously published.^34 35^ Expanded details are described in the Data Supplement.

### Electrophysiology

Kir2.1 currents were monitored in freshly isolated ECs and SMCs from third and fourth-order branches of mesenteric arteries using patch-clamp electrophysiology, as previously described.^36^ Expanded details are described in the Data Supplement.

### Oxidation-reduction production (ORP) measurements

Redox balance (integrated measure of the balance between total oxidants and reductants) was evaluated in plasma samples obtained from control and TBI rats by measuring the oxidation-reduction potential (ORP), or total oxidizing capacity^37^. Expanded details are described in the Data Supplement.

### PLA_2_ Activity Assay

Total PLA_2_ activity was measured in plasma samples obtained from control and TBI rats according to manufacturer’s instructions (BioVision, San Jose, CA). Briefly, fluorescence readings were taken with a microplate reader every 17 seconds for 45 min. Activity was calculated from the linear range of the reaction and corrected for volume and time.

### Pressure Myography

Pressure myography studies were conducted as previously reported^13^. Expanded details are described in the Data Supplement.

### Statistics

Metabolic pathway analysis, PLS-DA, heat mapping, and hierarchical clustering were performed using the MetaboAnalyst 3.0 package (www.metaboanalyst.com).^38^ Hierarchical clustering analysis (HCA) was also performed through the software GENE-E (Broad Institute, Cambridge, MA, USA). GraphPad Prism software (version 6.03; GraphPad Software, La Jolla, CA) was used for X-Y graphing and analysis; values are presented as means ± standard error of the mean. Differences were considered significant if *P*<0.05. Expanded details are described in the Data Supplement.

## Results

### TBI impairs vasoconstriction to Kir2.1 blockade and dilation to elevated extracellular potassium

To assess vascular Kir2.1 channel function, the inhibitors BaCl_2_ (100 μmol/L; Figure 1A) and ML 133 (20 μmol/L; Figure 1B) were exogenously applied to mesenteric arteries isolated from control and TBI rats. We found that vasoconstrictions to BaCl_2_ and ML 133 were both significantly decreased in TBI arteries when compared to controls. Next, a concentration response curve was performed by exchanging the arteriography chamber buffer with increasing steps of extracellular potassium (K^+^). Dilations to 10 mmol/L extracellular K+ were significantly reduced in MAs after TBI (Figure 1C). Concentrations at and greater than 14 mmol/L K^+^ constricted MAs in both groups. Interestingly, constrictions at the maximal concentration of 60 mmol/L K^+^ remained unaltered in TBI animals, suggesting preserved contractility following injury (Figure 1C). These findings suggest that vascular Kir2.1 channel function is crippled following TBI.

**Figure 1.**
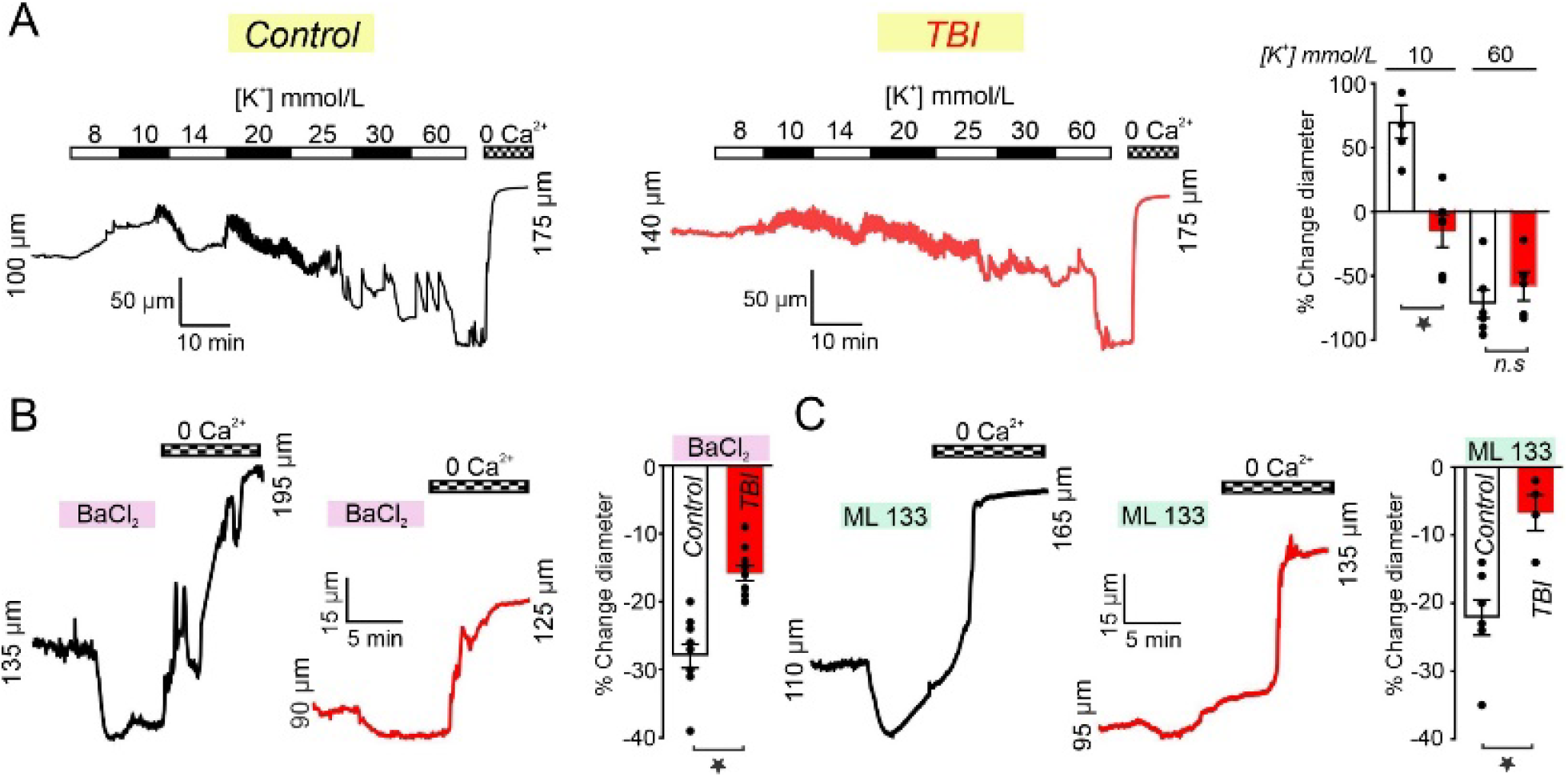
TBI impairs vasoconstrictions to the Kir2.1 channel blockers BaCl_2_ and ML 133 and dilations to extracellular K^+^ in mesenteric arteries. (A) Raising extracellular K^+^ from 6 to 8 mmol/L caused an immediate 43 ± 13 % (n=4) vasodilation and a subsequent peak dilation (62 ± 13 %; n=4) in control MAs pressurized to 80 mm Hg. As the concentration of extracellular K^+^ continued to increase from 14 to 60 mmol/L constrictions were observed in both control and TBI arteries. In MAs from TBI animals responses to 10 mmol/L K^+^ were significantly diminished when compared to controls (−15 ± 13 %; n=6; *P<0.05;* unpaired t-test). (B) BaCl_2_ (100 μmol/L)-induced constrictions were significantly diminished in TBI arteries (−16 ± 1.1 %; n=10) when compared to controls (−28 ± 1.7 %; n=12; *P<0.0001;* unpaired t-test). (C) Constrictions to ML 133 (20 μmol/L) were significantly reduced in TBI arteries (−6.8 % ± 2.6 %; n=4) when compared to controls (−22 ± 2.6 %; n=7; *P=0.0091;* unpaired t-test).

### Endothelial and smooth muscle Kir2.1 currents are significantly diminished following TBI and subsequently rescued by exogenous PIP_2_

We employed patch-clamp electrophysiology in native EC and SMCs from mesenteric arteries from TBI and control animals. First, we utilized a perforated patch-clamp configuration, in which the cytoplasm remains intact, to measure Kir2.1 currents in freshly isolated cells bathed in a high extracellular [K^+^]_o_ solution (60 mmol/L), used to amplify the inward component of Kir2.1 current amplitude. Under these conditions, the K^+^ equilibrium potential (E_*K*_) was −23 mV. We found in both EC and SMC barium-sensitive currents were significantly reduced following TBI (Figure 2A-B). One possible explanation for impaired Kir2.1 currents is the loss of the essential co-factor, PIP_2_. In support of this, we found the reduction in the Kir2.1 current observed in endothelial cells after trauma was partially restored when the synthetic PIP_2_ analog phosphatidylinositol 4,5-bisphosphate diC8 (10 μmol/L) was included in the intracellular pipette solution. Using conventional whole-cell configurations, we found current densities in the TBI cells treated with PIP2-diC8-were not significantly different from either the control PIP2-diC8-treated or naïve control groups (Figure 3).

**Figure 2.**
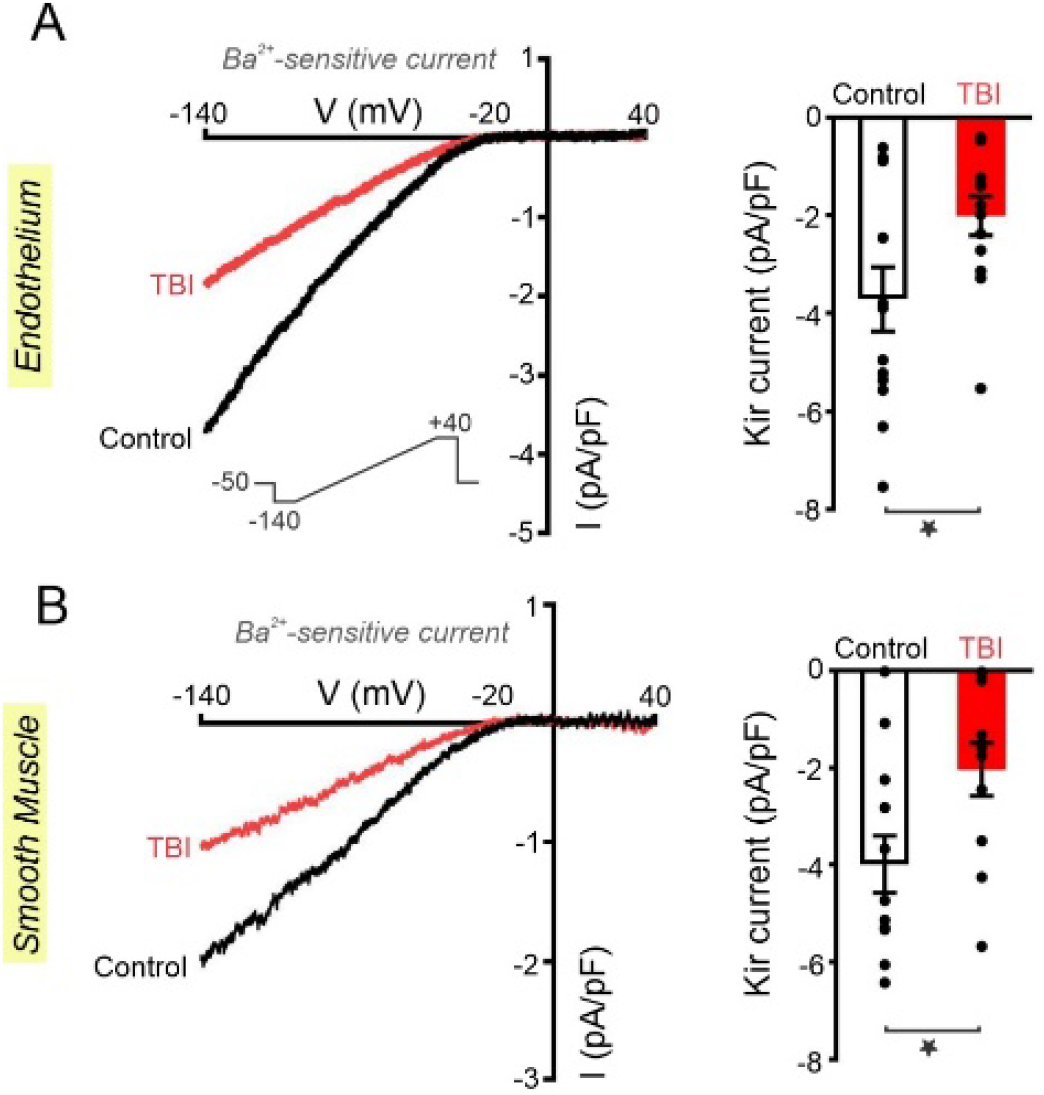
TBI diminishes Kir2.1 currents in mesenteric endothelial and smooth muscle cells. Whole-cell current was measured in freshly isolated endothelial and smooth muscle cells in a high extracellular K^+^ solution (60mmol/L) using a voltage ramp protocol (−140 to +40 mV) and perforated whole-cell patch-clamp configuration, in the absence and presence of BaCl_2_ (100 μmol/L) (A) Representative whole-cell recordings of Ba^2+^-sensitive currents in endothelial cells (ECs) from control (n=13) and TBI (n=13) rats. Kir2.1 current density at −140 mV is significantly diminished in ECs from TBI rats when compared to controls (−2.0 ± 0.4 pA/pF, n=13, vs. −3.7 ± 0.7 pA/pF, n=13, *P=0.03*, unpaired t-test). B) Representative whole-cell recording of Ba^2+^-sensitive currents in smooth muscle cells (SMCs) from control (n=12) and TBI (n=11) animals. Similarly, SMC Kir2.1 current density at −140 mV is significantly diminished after TBI when compared to controls (−1.01 ± 0.3 pA/pF, n=11, vs. −1.9 ± 0.3 pA/pF, n=12, *P=0.02*, unpaired t-test) (Figure 2B).

**Figure 3.**
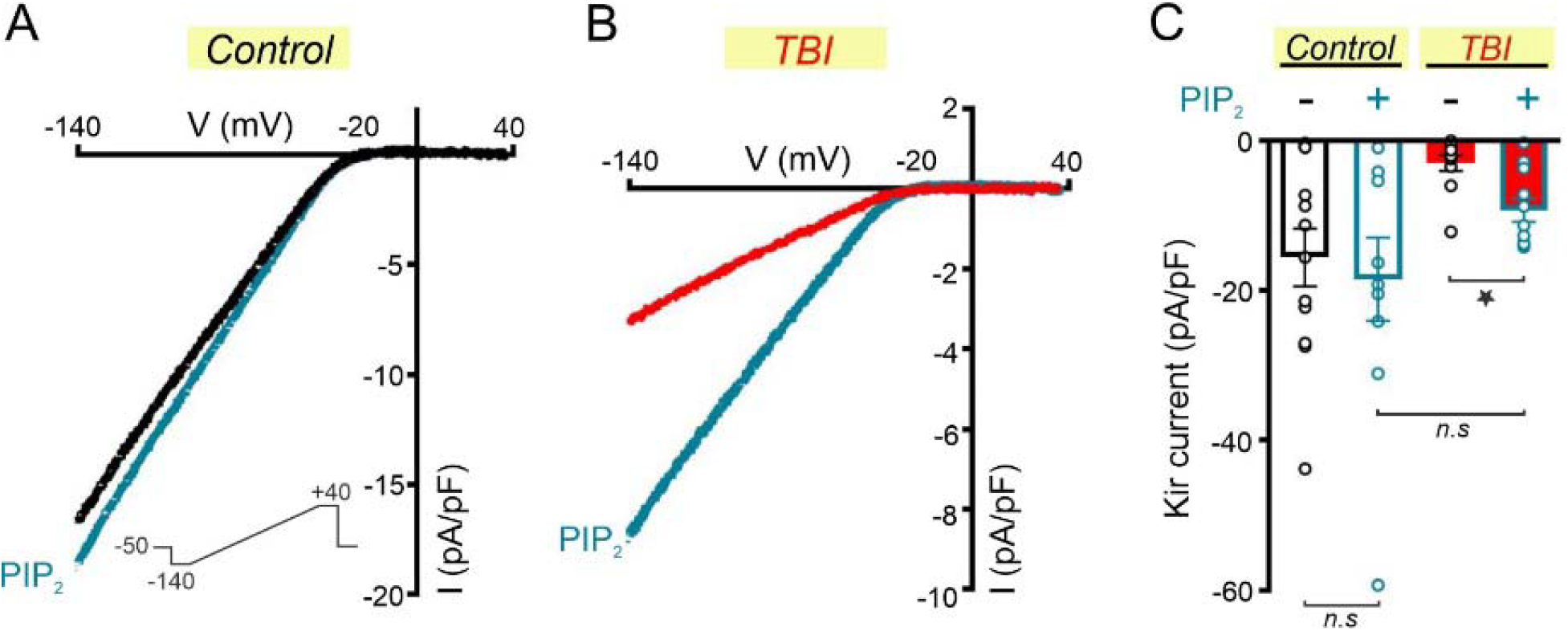
The PIP_2_ analog phosphatidylinositol 4,5-bisphosphate diC8 rescues endothelial Kir2.1 currents after TBI. Whole-cell currents were measured in freshly isolated endothelial and smooth muscle cells in a high extracellular K^+^ solution (60 mmol/L) using a voltage ramp protocol (−140 to +40 mV) and conventional whole-cell patch clamp configuration, in the absence and presence of BaCl_2_ (100 μmol/L). The PIP_2_ analog phosphatidylinositol 4,5-bisphosphate diC8 (10 μmol/L) was included into the patch pipette. (A) Representative wholecell recording of Ba^2+^-sensitive currents from control and TBI MAs in the absence or presence of PIP_2_-diC8 (10 μmol/L). (B) Summary data of peak 2 inward current (−140) in endothelial cells dialyzed with or without PIP_2_ from control or TBI groups. Kir2.1 current density was significantly diminished after TBI in MA ECs when compared to control ECs without PIP_2_ (−2.9 ± 1.0 pA/pF, n=11 vs. −15.7 ± 3.8 pA/pF, n=12; *P<0.0062;* unpaired t-test). No difference was observed in Kir2.1 current density (−140 mV) in the presence or absence of PIP_2_ in ECs from control MAs (−18.5 ± 5.5 pA/pF, n=10, vs. −15.7 ± 3.8 pA/pF, n=12; n.s.; unpaired t-test). Kir2.1 current density was significantly increased in TBI ECs in the presence of PIP_2_ when compared to TBI ECs without PIP_2_ (−8.6 ± 1.6 pA/pF, n=12, vs. −2.9 ± 1.0 pA/pF, n=11; *P<0.0078;* unpaired t-test). There was no significant difference in Kir2.1 current density in control and TBI ECs in the presence of PIP_2_, implying a rescue of functional Kir2.1 channels and a return to control-level current densities in the ECs from TBI MAs.

### TBI elevates hydrogen peroxide-derived oxidative stress and phospholipase A_2_ activity

We previously showed uncoupling of eNOS in systemic arteries after TBI; this would be expected to increase vascular H_2_O_2_ levels.^13^ Oxidative stress is known to activate PLA_2_ and contribute to the deranged lipidome observed after TBI.^39 40 41 42 43^ We therefore sought to measure global oxidant levels in plasma, and the response of mesenteric arteries to catalase, after TBI. We measured oxidation-reduction potential (ORP) in plasma before and after the addition of catalase (500 U/mL). The ORP after TBI was significantly higher compared to controls and was reduced to a level equal to controls by catalase, suggesting H_2_O_2_ is the specific reactive oxidant species present in TBI plasma (Figure 4A). In pressurized mesenteric arteries, constrictions to catalase (500 U/mL) were significantly increased following TBI (Figure 4B). In addition, total PLA_2_ activity (mU/mL) was significantly increased in plasma from injured animals when compared to controls (Figure 4C). These findings support a model of H_2_O_2_-derived PLA_2_ activity which would be expected to alter lipid metabolism and PIP_2_ levels after TBI.

**Figure 4.**
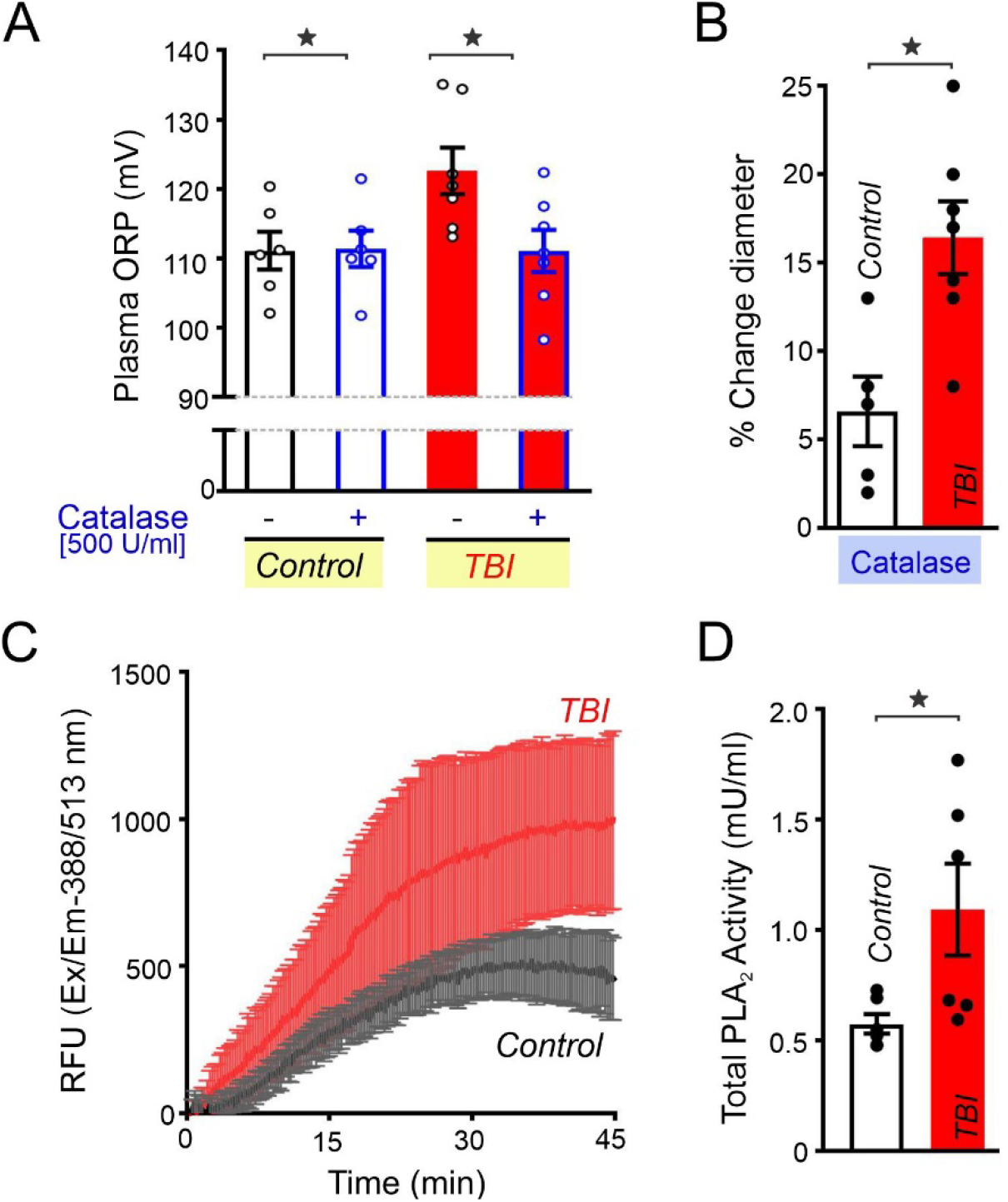
Hydrogen Peroxide (H_2_O_2_) levels and Phospholipase A2 (PLA_2_) activity are significantly increased in mesenteric arteries and plasma from TBI rats. (A) Oxidative reduction potential (ORP) measurements using the Redoxsys platform were conducted in paired plasma samples from control and TBI rats with and without the addition of catalase (500 U/mL). Plasma ORP was unchanged in control samples before and after the addition of catalase (500 U/mL) (111.1 ± 2.7 mV, vs. 111.4 ± 2.6 mV, n=6; n.s.; paired t-test). ORP in TBI plasma samples was significantly decreased to control values after the addition of catalase (500 U/mL) (122.6 ± 3.6 mV vs. 111.1 ± 3.0 mV, n=7, P=0.0017, Paired t-test). (B) Constrictions to catalase (500 U/mL), were significantly increased in mesenteric arteries from TBI rats when compared to controls (16.4 ± 2.01 %, n=7, vs. 6.6 ± 2.0 %, n=5, *P=0.0077;* unpaired t-test). (C) Averaged time course of PLA_2_ activity in control and TBI plasma samples. Total PLA_2_ activity is significantly increased in plasma from TBI rats when compared to controls (1.095 ± 0.21 mU/mL, n=6, vs. 0.576 ± 0.05 mU/mL, n=6, *P=0.0352;* unpaired t-test).

### TBI disrupts lipid metabolism causing accumulation of PIP_2_ degradation products

A high-throughput, semi-targeted metabolomics approach was applied to evaluate the metabolome of rats in our model of TBI, which has been shown to induce endothelial dysfunction in remote blood vessels from the mesenteric circulation. ^13^A correlation matrix and heat map for the top 50 metabolites with the lowest *P* values studied in plasma from control and TBI rats are shown (Figure 5A). Partial least squares discriminant analysis (PLS-DA) of control and TBI groups of metabolite levels in each biological replicate showed that the trauma and control samples clustered in distinct metabolic phenotypes (Figure 5B). Fold-change comparison between TBI and control animals are shown as graphs of lipid metabolites (Figure 5C). Of particular interest, we noted that TBI produced profound alterations in certain lipids, including increases in circulating eicosatetraenoic acid and AA, which can derive from PIP_2_ degradation (Figure 5C). Downstream products of AA metabolism were also altered, including prostanoids, eicosanoids, and leukotrienes (Figure 5C).

**Figure 5.**
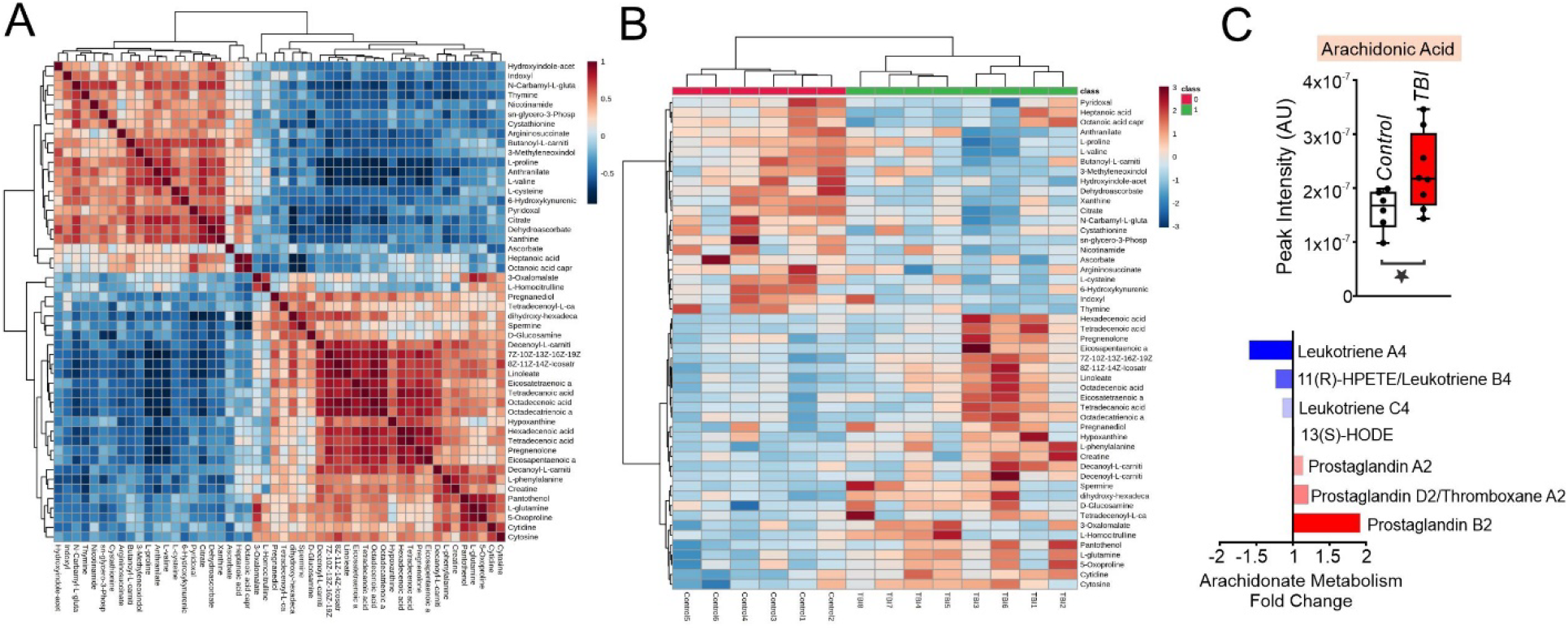
TBI causes accumulation of PIP_2_ degradation products and a profound lipidopathy. (A) Correlation matrix (distance measure: Pearson r) of the top 50 metabolites sorted by lowest *P* value and a heat map (distance measure: Euclidean; clustering algorithm: Ward) of the same data set. (B) Partial least squares discriminant analysis (PLS-DA) of control and TBI groups. Scores plot with Component 1 on the x-axis contributes to 40.1 % of the total variance. Component 2 on the y-axis contributes to 8.7 % of the total variance. (C) Circulating arachidonic acid (eicosatetraenoic acid) is significantly increased in plasma from TBI rats when compared to controls (2.3 x 10^7^ median AU, n=8, vs. 1.6 x 10^7^ median AU, n=6, *P=0.05*). Arachidonate metabolism pathway fold changes after TBI.

## Discussion

This study has three relevant findings. First, TBI cripples systemic vascular Kir2.1 channel function, evident in both single cell (EC and SMC) and intact vascular preparations. This has important clinical implications, as vascular Kir2.1 signaling modulates blood flow.^36^ Second, we provide a plausible mechanism for trauma-induced endothelial dysfunction; an increase in circulating and cellular H_2_O_2_-derived oxidative stress which causes an increase in PLA_2_ activity that would be expected to degrade PIP_2_ and thereby altering Kir2.1 channel function. Third, the fact that Kir2.1 channel function was rescued with PIP_2_ provides a potential therapeutic strategy that might improve endothelial function. Additionally, we deomonstrate that an isolated TBI, even without shock or hypoxia, produces a profound lipidopathy and an increase in H_2_O_2_ levels in both plasma and small blood vessels. A metabolomics screen was conducted to determine metabolic alterations in our TBI model which might impact vascular functions. This probe identified dysregulation of lipid metabolism pathways, including alteration of arachidonate pathways and phospholipid synthesis, and fatty acid mobilization/oxidation which are hallmarks of inflammation. Increases in H_2_O_2_-derived oxidative stress and phospholipase activity in plasma were consistent with upregulation of lipid metabolism.^42 39 18 44^ We focused on PIP_2_, a specific membrane phospholipid and key modulator of Kir2.1, the molecular feature that regulates myogenic tone and electrical conduction through the vascular endothelium.^45 46 47 48 27^

### Endotheliopathy of trauma

Recent studies have reported that biomarkers of endothelial dysfunction correlate with injury severity and adverse outcomes including coagulopathy and multi-organ dysfunction in trauma patients. Biomarkers reflecting endothelial damage or activation by cytokines include syndecan-1, thrombomodulin, and E-selectin.^8 10 7 49^ These biosignatures are elevated after isolated TBI and/or multi-system injury, in adults and children, and in both the prehospital setting and after resuscitation. These reports have suggested the existence of a pervasive “endotheliopathy of trauma” which complicates recovery and prevents optimizing the management of uncontrolled hemorrhage and coagulopathy; yet, few studies have actually evaluated endothelial cell or vascular function directly after trauma. Animal models of trauma provide the opportunity to directly study blood vessels to further elucidate the cellular mechanisms of endothelial injury.

We previously reported that after TBI, blood vessels harvested from the mesenteric circulation, remote from the site of injury, showed impaired endothelial-dependent vasodilatory function, but Kir2.1 channels were not directly assessed.^13^ Here, we show evidence of systemic endothelial dysfunction after TBI, showing that Kir2.1 responses of pressurized arteries and in isolated vascular cells are significantly diminished, providing a novel model to explain remote endothelial effects of severe trauma.

### Kir2.1 channel dysfunction and PIP_2_ regulation

The Kir2.1 channel is the molecular cornerstone responsible for dynamic regulation of blood flow in the brain and other excitable tissues, and the membrane phospholipid PIP_2_ is required for channel function.^20^ Kir2.1 channels are exquisitely sensitive to changes in the gradient of extracellular K^+^, which acts as a potent electrochemical vasodilator. In the brain, capillary endothelial Kir2.1 channels play a critical role in the dynamic regulation of regional perfusion, by conducting a spreading hyperpolarizing signal in response to locally elevated extracellular K^+^ released during neural activity, and thereby directing blood flow^27^. We recently demonstrated that brain capillary endothelial Kir2.1 possess the ability to respond to K^+^ released during neuronal activity causing a hyperpolarizing electrical signal that propagates in a retrograde fashion to the parenchymal arterioles.^48^ Cerebral ischemia leads to a loss of functional Kir2.1 channels in parenchymal arteriole smooth muscle, impairing neurovascular coupling.^50^ We recently found that a TBI results in impaired Kir2.1 channel function in capillary ECs from the opposite side of the brain, and we hypothesized that this dysfunction may be widespread and not limited to cerebral endothelium. Endothelial Kir2.1 channels are also important in mesenteric resistance arteries, boosting vasodilatory signals through endothelial derived hyperpolarizing pathways.^45^ Our results support a model by which systemic oxidative stress, involving H_2_O_2_, acts via PLA_2_ to metabolize PIP_2_ and therefore disrupt Kir2.1 function (Fig. 6). Our model provides a plausible mechanism by which the metabolic stress of trauma leads to systemic endothelial dysfunction, based on the known link between H_2_O_2_ and PLA_2_ activation.^39 51 52 44^

**Figure 6.**
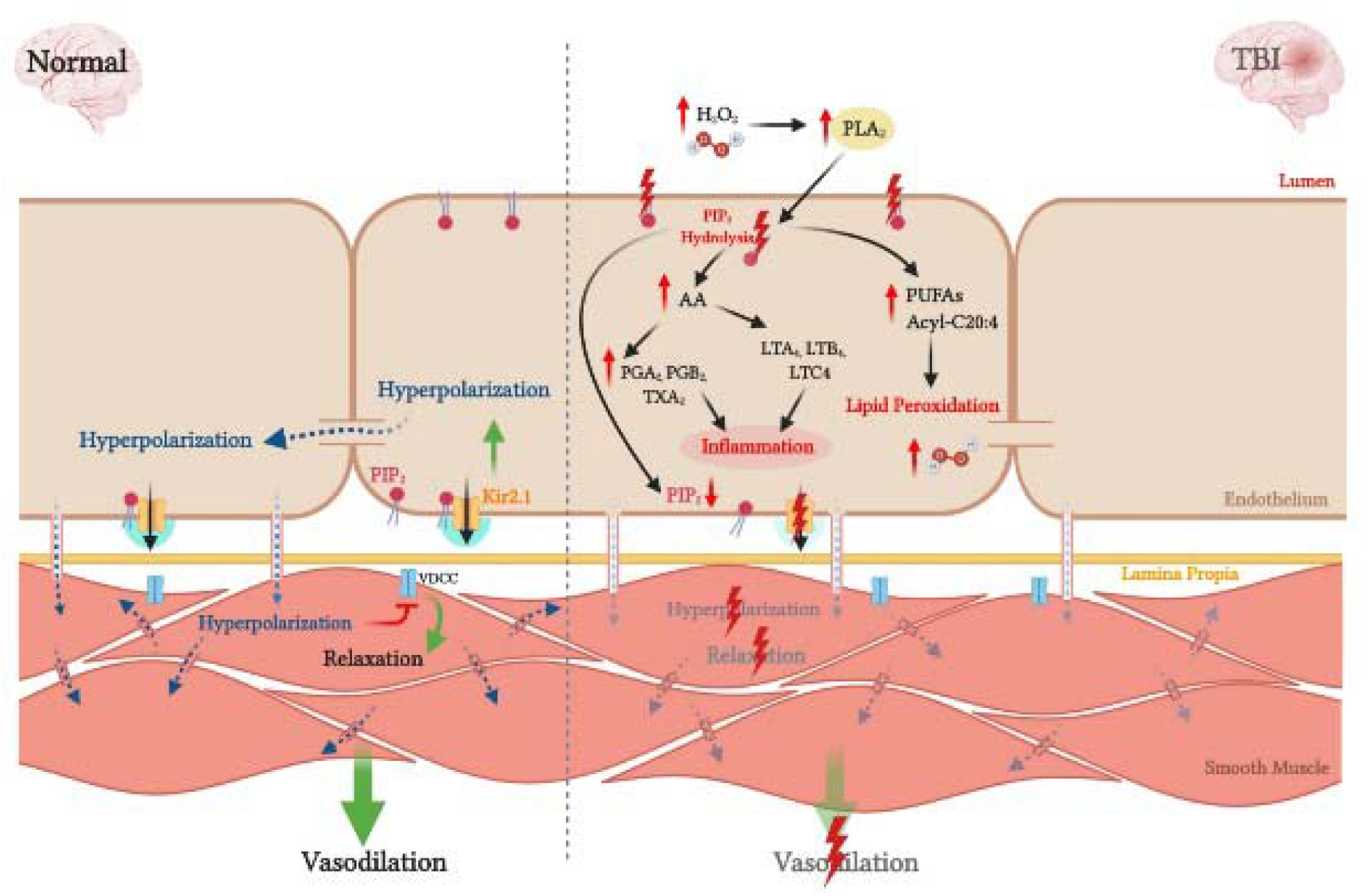
Schematic Overview. As PIP_2_ is required to stabilize Kir2.1 channel activity, and hydrogen peroxide (H_2_O_2_) drives down PIP_2_ levels, we hypothesized that TBI results in impaired Kir2.1 channel function in MAs. We demonstrated that 24 hours following TBI an increase in circulating and cellular H_2_O_2_-derived oxidative stress causes an increase in PLA_2_ activity which degrades PIP_2_ in the cell membrane, thereby impairing Kir2.1 channel function and subsequent hyperpolarization/vessel dilation.

### Deranged lipid metabolism, H_2_O_2_ and PLA_2_ after TBI

Providing further support for our model, our metabolomics screen demonstrated that lipid metabolism homeostasis is significantly deranged as early as one day after TBI. These findings correlate with prior reports on TBI during the sub-acute and chronic phases suggesting that disturbed lipid metabolism is a strong predictor of clinical outcome.^31 17^ With the aid of mass spectrometry-based lipidomics, they provided quantitative characterization of lipid peroxidation products after TBI and acknowledge PLA_2_ as an important mediator in the hydrolysis of peroxidized phospholipids.^52^ The polyunsaturated fatty acid component of membrane phospholipids are often targets for peroxidation, broadly defined as the process of inserting a hydroperoxy group into a lipid. Peroxidation increases the rates of hydrolysis by phospholipases. The extent of lipid peroxidation after severe traumatic brain injury correlates with injury severity and mortality.^52^

Rodent models have also provided insight into specific changes in lipid metabolism induced by trauma. Major changes in free fatty acids, including docosahexaenoic, stearic, oleic, and arachidonic acids, were sustained from 4 to 35 days after injury in a rat.^53^ In addition, significant upregulation in polyunsaturated fatty acids (PUFAs) and PUFA-containing diacylglycerols and changes in membrane phospholipids including sphingolipids have been observed at one week after injury.^16^ Lipidomic analyses in mouse model of TBI identified injury-specific phospholipid changes that persisted as long as 3 months after trauma induction.^14^ Our findings showed that not only lipid metabolites were altered after TBI, but also, plasma H_2_O_2_ and PLA_2_ activity were significantly increased in these animals. Specific metabolites of PLA_2_-dependent PIP_2_ degradation products, such as arachidonic acid and its downstream metabolites, were effectively detected in our model of TBI. In this context, increased oxidative stress has been described after severe trauma and in TBI in particular.^54^

## Conclusions

Collectively, our results support a novel mechanism to explain remote endothelial effects of severe for systemic endothelial dysfunction after TBI, through diminished Kir2.1 responses which can be rescued by PIP_2_. These findings may be generalizable to other pathological conditions characterized by altered lipid metabolism pathways and reductions in PIP_2_ levels, and suggest PIP_2_ may be a potential therapeutic target to improve endothelial function in conditions characterized by altered lipid metabolism such as brain trauma and stroke.

## Non-standard Abbreviations and Acronyms

TBI: Traumatic Brain Injury
Kir: Inward Rectifier Potassium Channel
SMC: Smooth Muscle Cell
EC: Endothelial Cell
MA: Mesenteric Artery
PIP_2_: Phosphatidylinositol 4,5-bisphosphate
PLA_2_: Phospholipase A_2_
H_2_O_2_: Hydrogen Peroxide
AA: Arachidonic Acid

## Acknowledgements

A.M.S. designed experiments, acquired and analyzed data, wrote the initial draft and edited all subsequent drafts of the manuscript. N.V. assisted with experimental design and participated in all stages of critical revisions. A.B. acquired and analyzed electrophysiology data. A.D. and T.M. acquired and analyzed mass spectrometry data. M.S. contributed to data representation and in critical revisions of the manuscript. O.H. participated in critical revisions of the manuscript. M.T.N. and K.F. directed the study and edited the manuscript. All authors reviewed the manuscript, contributed to critical revisions, and approved its submission.

## Sources of Funding

Department of Defense / The Henry M. Jackson Foundation for the Advancement of Military Medicine (HU001-18-2-0016 to KF and MTN), NIH (R01GM123010 to KF; UM1 HL120877, R01NS110656, and R35HL140027 to MTN; P20-GM-135007 to MTN and OFH), EC Horizon 2020 (MTN), Foundation Leducq (MTN), and the Totman Medical Research Trust (MTN).

## Conflicts-of-Interests/Disclosure(s)

### Disclosures

O.F.H. and M.T.N. are inventors of patent number 62/823,378 “Methods to promote cerebral blood flow in the brain” which was submitted on 3/25/2019.

- **Expanded Materials & Methods**
- **References 13; 29; 30; 36**

