## Supplemental Methods for "Traumatic Brain Injury Impairs Systemic Vascular Function Through Altered Lipid Metabolism and Disruption of Inward-Rectifier Potassium (Kir2.1) Channels"

**SUPPLEMENTAL MATERIAL**

***Animal and injury model****.* Animals were anesthetized with inhaled 2 to 5 % isoflurane prior to TBI or sham surgery. All animals received buprenorphine analgesia (subcutaneous; 0.05 mg/kg) while under anesthesia and at 6 to 12 hours after surgery. Briefly, a primary injury was induced by a direct contusion to the brain delivered to the left cerebral hemisphere by a pendulum impacting a fluid-filled chamber connected to the intact dura through a craniotomy.^29 30^ A fluid percussion injury was induced to a target pressure of < 482 kPa (< 70 PSI) over a 500 msec period; the pressure was transduced and measured in each surgery. This level allows > 90% recovery, defined as ability to maintain upright posture, ambulate, and take oral hydration, and produces a highly reproducible outcome of moderate brain injury severity in those surviving animals. These experimental animals have measurable deficits in sensorimotor coordination, along with significant cardiovascular and cerebrovascular effects.^29 30^ Control animals were subjected to scalp incisions but without the percussion injury. At 24 h after recovery from surgery, animals were euthanized using deep pentobarbital anesthesia (intraperitoneal; 0.03 mg/kg) and tissues were collected for experiments.

***Sample Preparation*** ***and UHPLC-MS analysis.*** Prior to LC-MS analysis, samples were placed on ice and diluted with 24 volumes of methanol:acetonitrile:water (5:3:2, v:v). Suspensions were vortexed continuously for 30 min at 4°C. Insoluble material was removed by centrifugation at 10,000 g for 10 min at 4°C and supernatants were isolated for metabolomics analysis by UHPLC-MS. The analytical platform employs a Vanquish UHPLC system (Thermo Fisher Scientific, San Jose, CA, USA) coupled online to a Q Exactive mass spectrometer (Thermo Fisher Scientific, San Jose, CA, USA). Samples were resolved over a Kinetex C18 column, 2.1 x 150 mm, 1.7 µm particle size (Phenomenex, Torrance, CA, USA) equipped with a guard column (SecurityGuard^TM^ Ultracartridge – UHPLC C18 for 2.1 mm ID Columns – AJO-8782 – Phenomenex, Torrance, CA, USA) using an aqueous phase (A) of water and 0.1% formic acid and a mobile phase (B) of acetonitrile and 0.1% formic acid for positive ion polarity mode, and an aqueous phase (A) of water:acetonitrile (95:5) with 1 mmol/L ammonium acetate and a mobile phase (B) of acetonitrile:water (95:5) with 1 mmol/L ammonium acetate for negative ion polarity mode. Samples were eluted from the column using either an isocratic elution of 5% B flowed at 250 µl/min and 25ºC or a gradient from 5% to 95% B over 1 minute, followed by an isocratic hold at 95% B for 2 minutes, flowed at 400 µl/min and 45ºC. The Q Exactive mass spectrometer (Thermo Fisher Scientific, San Jose, CA, USA) was operated independently in positive or negative ion mode, scanning in Full MS mode (2 μscans) from 60 to 900 m/z at 70,000 resolution, with 4 kV spray voltage, 45 shealth gas, 15 auxiliary gas. Calibration was performed prior to analysis using the Pierce^TM^ Positive and Negative Ion Calibration Solutions (Thermo Fisher Scientific. Metabolite assignments, isotopologue distributions, and correction for expected natural abundances of deuterium, ^13^C, and ^15^N isotopes were performed using MAVEN (Princeton, NJ, USA). ^55^

***Electrophysiology.*** Briefly, mesenteric arteries were harvested and endothelial cells were isolated in 1.5 mL dissociation solution containing (in mmol/L): 134 KCl, 6 KOH, 10 NaOH, 1.1 MgCl_2_, 1.8 CaCl_2_, 5 EGTA, 10 HEPES (pH 7.3), containing 0.5 mg/mL neutral protease and elastase (0.5 mg/mL; Worthington Biochemical Corp., Lakewood, NJ), for 60 minutes at 37°C. Collagenase (0.5 mg/mL; Worthington Type I) was added to the dissociation solution for three more minutes. Vessels were then washed in Ca^2+^-free dissociation solution (4°C) for 5 to 10 minutes and the solution triturated 10 times through a custom glass Pasteur pipette to release single endothelial cells into the solution. Smooth muscle cells were isolated in dissociation solution containing 1 mg/mL papain, 0.5 mg/mL dithioerythritol (DTE), and 0.5 mg/mL bovine serum albumin (BSA) for 25 minutes. Next, 1 mg/mL of collagenase (Worthington Type IV), 0.25 mg/ml elastase, and 0.5 mg/mL Trypsin inhibitor and 100 µM CaCl_2_ were exchanged in the solution for 10 min. Electrophysiology was performed in either the conventional or perforated whole-cell configuration. Whole-cell currents were amplified using an Axopatch 200B amplifier (Molecular Devices, USA), filtered at 1 kHz, digitized at 10 kHz, and stored on a computer for offline analysis with Clampfit 10.7 software. Patch pipettes were pulled from borosilicate, microcapillary tubes (1.5-mm O.D., 1.17-mm I.D.; Sutter Instruments, Novato, CA), and fire-polished (resistance ~ 4-6 MΩ). Cells were voltage-clamped at a holding V_M_ of -50 mV and equilibrated for 15 minutes in a bath solution containing (in mmol/L): 134 NaCl, 6 KCl, 1 MgCl_2_, 10 glucose, 2 CaCl_2_, and 10 HEPES (pH 7.4). For the perforated-patch configuration, pipettes were backfilled with a solution containing (in mmol/L): 10 NaCl, 30 KCl, 110 K^+^-Aspartate, 1 MgCl_2_, 10 HEPES (pH 7.2), and 200 to 250 µg/mL amphotericin B, added freshly on the day of the experiment. For the conventional whole-cell configuration, the pipette solution was composed of (in mmol/L): 134 KCl, 6 KOH, 10 NaOH, 1.1 MgCl_2_, 1.8 CaCl_2_, 5 EGTA, 10 HEPES (pH 7.2). Ba^2+^-sensitive Kir2 currents were quantified by elevating extracellular [K^+^] from 6 to 60 mmol/L via equimolar replacement of NaCl by KCl.A 400-ms voltage-ramp protocol (-140 mV to +40 mV) was applied. All experiments were performed at room temperature (~ 22°C). Control EC capacitance was 10.7 ± 0.5 pF and TBI EC capacitance was 11.0 ± 0.6 pF. Control SMC capacitance was 11.2 ± 1.0 pF and TBI SMC capacitance was 14.1 ± 0.7 pF.

***Oxidation-reduction production (ORP) measurements***. Whole blood was collected at the time of euthanasia into an evacuated tube containing heparin and immediately centrifuged (2,000 rpm; 4°C). Plasma samples (30 µL) were collected and tested using the RedoxSYS diagnostic platform, consisting of a micro Pt/AgCl combination redox electrode sensor and benchtop analyzer (Aytu Bioscience, Inc., Englewood, CO). Values were recorded in mV after ORP readings were stable for 10 sec. The diagnostic platform was calibrated before use and validated in a previous study. ^13^

***Pressure Myography.*** Immediately after euthanasia, a midline laparotomy was performed and the mesentery was dissected out and placed into cold (4°C) physiological saline solution (PSS) with the following composition (in mmol/L): 119 NaCl, 45 KCl, 24 NaHCO_3_, 1 KH_2_PO_4_, 2.5 CaCl_2_, 1 MgCl_2_, and 11 glucose (pH 7.4). The mesentery was then pinned out on a dissecting dish and fourth- and fifth-order mesenteric arteries were dissected free from the surrounding adipose and connective tissue. For each experiment, an individual mesenteric artery was cannulated in a pressure myograph (Living Systems Instrumentation, St. Albans, VT) containing oxygenated 20% O_2_/ 5% CO_2_ PSS at 37°C. Intraluminal pressure during the experiment was maintained at 80 mm Hg using a pressure servo system, and blood vessel diameters were measured using edge-detection software coupled to a camera (IonOptix, Westwood, MA). Arteries were equilibrated for 10 min, pressurized and allowed to develop spontaneous myogenic tone over the course of 30 min, defined as >20% constriction after equilibration and pressurization of vessel. The vascular endothelium was considered intact if a dilation of greater than 85% was elicited using the endothelial-dependent vasodilator NS309 (1 µM; Cayman Chemical Company, Ann Arbor, MI). Arteries which did not develop spontaneous myogenic tone were excluded.

***Statistics.*** Data was tested for normality and a parametric or non-parametric statistical test was subsequently applied. One- or two-way analysis of variance (ANOVA) was used for comparisons of multiple group measurements. In a few experimental series, a single control group was used to test multiple hypotheses. To avoid increasing the likelihood of a Type I error, we used the Bonferroni correction to test each individual hypothesis at a significance level determined by α = 1/m, where m is the number of comparisons.
